# Lowe Syndrome-linked endocytic adaptors direct membrane cycling kinetics with OCRL in *Dictyostelium discoideum*

**DOI:** 10.1101/616664

**Authors:** Alexandre Luscher, Florian Fröhlich, Caroline Barisch, Clare Littlewood, Joe Metcalfe, Florence Leuba, Anita Palma, Michelle Pirruccello, Gianni Cesareni, Massimiliano Stagi, Tobias C. Walther, Thierry Soldati, Pietro De Camilli, Laura E. Swan

## Abstract

Mutations of the inositol 5-phosphatase OCRL cause Lowe Syndrome (LS), characterized by congenital cataract, low IQ and defective kidney proximal tubule resorption. A key subset of LS mutants abolishes OCRL’s interactions with endocytic adaptors containing F&H peptide motifs. Converging unbiased methods examining human peptides and the unicellular phagocytic organism *Dictyostelium discoideum*, reveal that, like OCRL, the *Dictyostelium* OCRL orthologue Dd5P4 binds two proteins closely related to the F&H proteins APPL1 and Ses1/2 (also referred to as IPIP27A/B). In addition, a novel conserved F&H interactor was identified, GxcU (in *Dictyostelium)* and the Cdc42-GEF Frabin (in human cells). Examining these proteins in *Dictyostelium discoideum*, we find that, like OCRL, Dd5P4 acts at well-conserved and physically distinct endocytic stations. Dd5P4 functions in coordination with F&H proteins to control membrane deformation at multiple stages of endocytosis, and suppresses GxcU-mediated activity during fluid-phase micropinocytosis. We also reveal that OCRL/Dd5P4 acts at the contractile vacuole, an exocytic osmoregulatory organelle. We propose F&H peptide-containing proteins may be key modifiers of LS phenotypes.

## Introduction

Phosphoinositide lipids (PIPs) are a group of seven phospholipids which, by reversible phosphorylation of the 3’, 4’ and 5’ positions of their cytosolic inositol ring, help specify membrane function and identity. They achieve this chiefly by recruiting cytosolic proteins, such as adaptor proteins for membrane sorting and deformation complexes, allowing co-ordinated recruitment and sorting of transmembrane cargoes in the exo/endocytic cycle. As membranes progress through the exo/endocytic pathways, changes in their PIP “signature” signal a change in their identity, thus modifying membrane function. Humans express at least nine enzymes that act on the 5’ position of the lipids PI(4,5)P_2_ and/or PI(3,4,5)P_3_. Both OCRL (OCRL1) and its paralog INPP5B (also called OCRL2) contain an N-terminal PH domain ^1^, a central 5’-phosphatase catalytic module, an ASH (<u>A</u>SPM, <u>S</u>PD-2, <u>H</u>ydin) domain ^2^ which stabilises a C-terminal catalytically inactive RhoGAP-like domain^3,4^. The core structure of the OCRL proteins (5’ phosphatase and ASH-RhoGAP tandem domain) is exceptionally well conserved during evolution. OCRL-like proteins are found in most eukaryotic lineages, including distantly related eukaryotes such as the protist *Giardia lamblia*^5^, suggesting a significant conserved function.

OCRL acts on the 5-phosphates of PI(4,5)P_2_ and PI(3,4,5)P_3_, two PIPs which are typically enriched at the plasma membrane. However, OCRL is a promiscuous interactor of endocytic proteins, which are found on organelles not typically enriched in these substrate lipids. Endocytic interactors include proteins involved in the initial steps of clathrin-mediated endocytosis ^1,6-8^, early and late endosomal adaptor proteins^3,4,9,10^, a wide variety of Rab^11,12^ and Rho family GTPases^13^, leading to a distinctive distribution of OCRL-positive intracellular compartments ^3,7,14^.

Loss, truncation or missense mutation of OCRL causes the congenital X-linked disorder, oculo-cerebral renal syndrome of Lowe (Lowe Syndrome (LS; OMIM: 309000))^15^, affecting the brain, eyes and renal system, or a less severe condition that primarily involves later-onset dysfunction of the renal system, called Dent2 disease (OMIM: 300555)^16^. LS and Dent2 disease are caused by a similar spectrum of OCRL ^15^ mutations, including in one family, the same mutation ^17^. Missense mutations of the ASH-RhoGAP domain provoke the full spectrum of LS symptoms, namely profound defects in PI(4,5)P_2_ metabolism in patient fibroblasts and the clinical triad of kidney, brain and ocular disorders. Previously we have identified that these ASH-RhoGAP mutants are specifically deficient in binding to endocytic proteins containing a helical peptide that we called the F&H motif (for the obligate presence of conserved phenylalanine and histidine residues^4,10^).

The F&H peptide directly binds a surface unique to the OCRL/INPP5B RhoGAP domain. Previous to this study, we had identified F&H motifs in three endocytic proteins, the very early endocytic protein APPL1^3^, which is a resident of non-canonical PI3P-negative, Rab5-positive endosomes^18^ and a pair of later, endosomal adaptors Ses1 and Ses2^10^, otherwise known as IPIP27A/B^19^, which are resident on canonical PI3P-positive endosomes. Binding of APPL1 and Ses1/2 to OCRL is mutually exclusive and occurs at different endosomal stations defined by the absence or presence of PI3P respectively ^10^, suggesting formation of OCRL-F&H adaptor protein complexes is subject to regulation by other endocytic factors.

The function of F&H motif-dependent interactions of OCRL and INPP5B and their contribution to LS pathology remain elusive. In different systems, fluid-phase endocytosis ^20^ and efficient ciliary traffic^21^ require the interaction of OCRL with F&H proteins, suggesting their relevance to LS phenotypes. Ses2/IPIP27B recruits OCRL to sort cation-independent mannose-6-phosphate receptor (Ci-M6PR)^22^, a retromer cargo, demonstrating a requirement for OCRL-driven PI(4,5)P_2_ dephosphorylation *via* F&H proteins in late endocytic traffic. The F&H binding patch is conserved in most species that express an OCRL-like protein, including a variety of protists and other unicellular eukaryotes including *D. discoideum, Trypanosoma brucei*^4^, *Albugo candida* (CCI48278.1, an Oomycete), and *Cyanidioschyzon merolae* (XP_005535444.1, a red alga), suggesting that interactions with F&H peptides are critical to OCRL function. In the above organisms no orthologues of APPL1 and Ses1/2 could be identified by primary amino acid sequence similarity or domain structure predictions. We hypothesised that critical functions of OCRL and INPP5B were mediated by yet-unknown F&H peptide-containing binding partners that may have human orthologues, which may help to explain cellular dysfunction in LS.

This study aimed to identify ancestral F&H interactors to gain functional insight into the F&H peptide-binding interface of OCRL/INPP5B, and to help understand OCRL dysfunction in LS/Dent2 disease. We utilized the unicellular organism *D. discoideum*, where the presence of a single OCRL/INPP5B orthologue (Dd5P4, which has a predicted F&H-motif-binding surface) - avoids complications of vertebrate models that express two, partially redundant, enzymes^23,24^, OCRL and INPP5B. Importantly, we expected that the study of Dd5P4 would allow us to assess the role of F&H domain binding to OCRL/Dd5P4 in a cellular context where orthologues of APPL1 and Ses1/2 did not appear to be present.

Strikingly, our studies showed that not only a network of F&H peptide interactors of OCRL/INPP5B, but also the subcellular localization of these interactors are strongly conserved, suggesting that OCRL-driven hydrolysis of PI(4,5)P_2_ or PI(3,4,5)P_3_ on endolysosomal organelles is a widely conserved feature of membrane trafficking. We isolated three *D. discoideum* proteins with a *bona fide* F&H motif. Two are related in domain structure to APPL1 and Ses1/2, in spite of divergent amino acid sequence: an APPL1-like early endocytic BAR domain protein and a Ses1/2-like PH domain protein. A third interactor with a novel domain structure is a Rho family GEF, GxcU, orthologous to mammalian Frabin, which we find also has a conserved F&H motif, suggesting its relevance to LS/Dent pathology. We find that overexpression of GxcU leads to endocytic defects which are enhanced by Dd5P4 loss, suggesting Dd5P4 represses GxcU activity. We additionally reveal that both Dd5P4 and the APPL1-like protein are recruited to the contractile vacuole membrane of *D. discoideum* upon their kiss-and-run exocytic water discharge, providing new evidence for a role of OCRL-APPL1 partnership in membrane remodelling in the early endocytic pathway.

## Results

### Peptide array experiments confirm F&H motif and identifies Frabin/FGD4 as a potential F&H motif containing OCRL interactor

To search for potential new F&H motif-interactors of human OCRL, we first defined the F&H interaction consensus via peptide microarray experiments, to then search for proteins harbouring this consensus in the human proteome (**Figure 1** and **Supplementary Figure 1**). Each residue of the F&H peptide of human Ses1 (13 amino acids) was systematically substituted, and immobilised on nitrocellulose membranes. These membranes were then probed in an overlay assay with a GST fusion of the ASHRhoGAP domain of human OCRL (**Figure 1A**), or of the same domain harbouring a tryptophan to alanine mutation at amino acid 739 (i.e. a tryptophan critically required for F&H motif binding) ^4^ to exclude nonspecific interactors. We found that this assay validated the F&H motif deduced by evolutionary conservation of residues in APPL1 and Ses1/2 ^10^ (**Figure 1B,C**). Interaction with OCRL in the overlay assay was absolutely dependent on the presence of F and H residues at positions 2 and 6 of the motif as previously predicted. The peptide array assay, however, appears to favour an APPL1-like mode of binding^3,4^, which requires only 11aa of the 13aa F&H helical peptide (**Figure 1A,C,D**). The array-derived motif also includes either an R or hydrophobic residue as opposed to the bulky hydrophobic residue at position 9 defined by conservation (**Figure 1B**), generating a slightly less selective F&H motif than that derived by evolutionary conservation.

**Figure 1:**
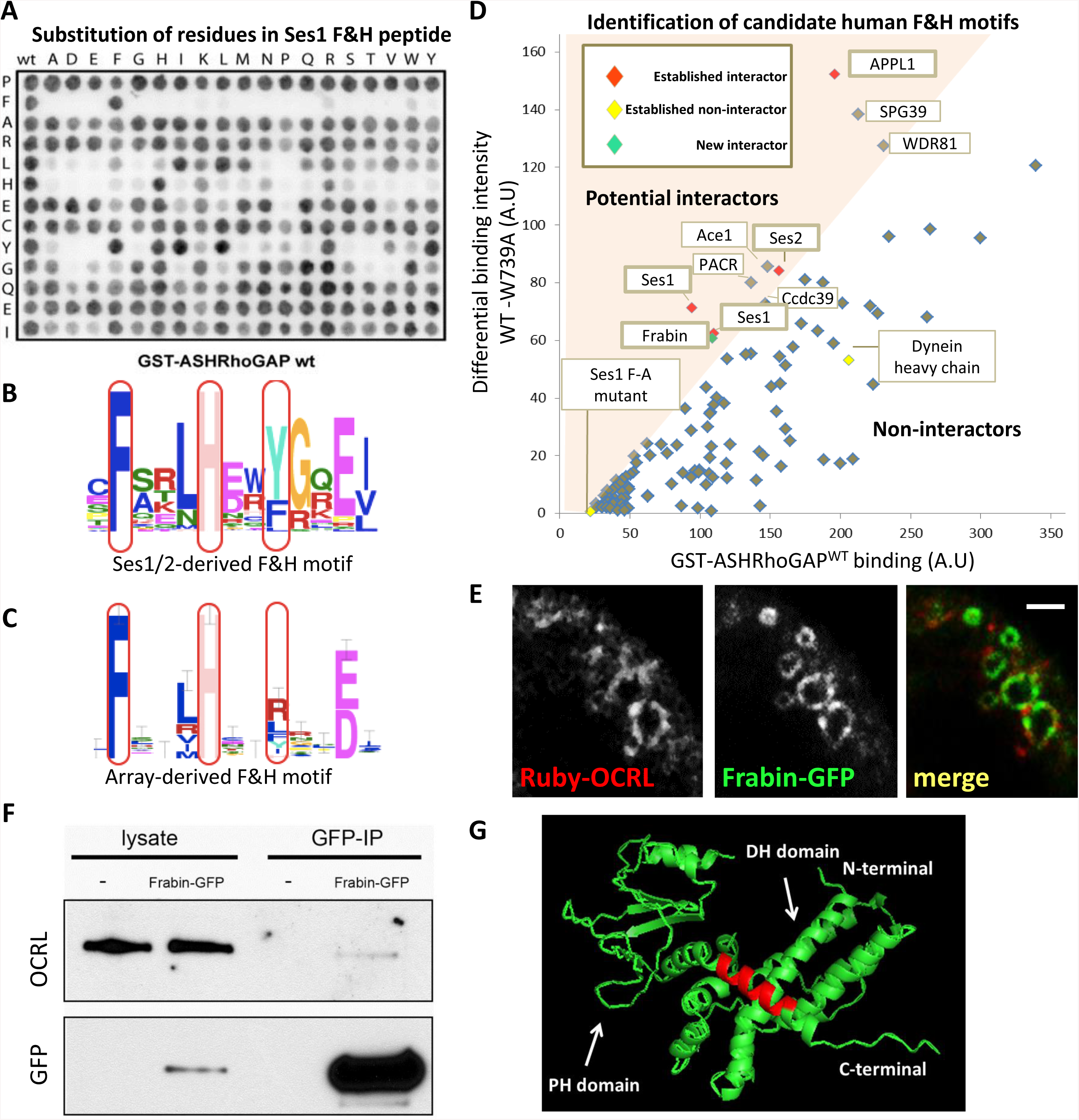
Overlay assay detects new F&H candidate proteins**. A)** Peptide overlay experiments confirm the established consensus sequence for F&H peptides. Each residue was systematically substituted in the Ses1 F&H peptide (*vertical*: the original residue in the peptide, *horizontal*: substituted residue) immobilised on nitrocellulose and probed with GST-hOCRL-ASHRhoGAP or the non-F&H binding GST-hOCRL-ASHRhoGAP^W739A^ to control for non-specific binding. **B**) F&H consensus derived from sequence conservation of F&H peptides in Ses1/2 proteins. **C**) F&H consensus derived from peptide overlay. **D**) Naturally occurring human peptides which fall within the F&H consensus derived in **C** were immobilised on nitrocellulose and probed with wild-type GST-hOCRL-ASHRhoGAP (**Figure 1D**, *x-axis*) and control as before. WT binding intensity is plotted against the difference (*y-axis*, arbitrary units) in intensity of binding to WT and W739A control. We defined candidate interactors as peptides in the upper left quadrant of the graph (*i.e* peptides whose binding to GST-hOCRL-ASHRhoGAP is reduced by over 50% by the W739A mutation). All known F&H peptide interactors (red) were isolated by this analysis, including two identical Ses1 peptides. Peptides determined as non-interactors by isothermal titration calorimetry are excluded (yellow) (**Supplementary Figure 1B**). The Frabin F&H peptide (green) was isolated as an interactor in this assay. **E**) Ruby-OCRL and Frabin-GFP are found together on pinosomal compartments in Cos7, see **Supplementary Movie 1**. Scale bar: 3μm. **F**) Full length Frabin-GFP immunoprecipitations isolated a weak but specific interaction with endogenous OCRL in Cos7 cells. **G**) predicted structure of the Frabin DH-PH module shows the Frabin F&H peptide (red) is found on an extended alpha helix of the Frabin PH1 domain.

Peptides in the human proteome that fit the peptide-array F&H consensus were synthesised and immobilised on nitrocellulose membranes and probed as above with GST-OCRL-ASHRhoGAP and GST-OCRL-ASHRhoGAP^W739A^ to isolate candidate F&H proteins in humans. Peptides predicted to be inaccessible to cytosolic proteins (i.e. peptides known to be buried in folded domains, transmembrane regions or extracellular domains) were excluded from the analysis. Peptides whose binding to GST-ASHRhoGAP is diminished by at least 50% by W739A mutation (coloured region in **Figure 1D**) were considered to be candidate F&H peptides.

This screen identified known F&H peptide interactors Ses1, Ses2, APPL1, as expected, and excluded peptides that we had previously determined using isothermal titration calorimetry (ITC) to be non-specific^4,10^, (**Supplementary Figure 1**) confirming the specificity of the analysis. It also identified additional potential interactors: PACR, Ace1, Ccdc39, SPG39, WDR81 and Frabin (**Figure 1D**). PACR was excluded on closer inspection as key residues are buried in the plasma membrane^25^. Ace1, Ccdc39 and SPG39 were also excluded as, based on western blotting on GST-ASHRhoGAP pulldowns in HKC cells (**Supplementary Figure 1**) full-length proteins did not show a specific interaction with the F&H interaction surface of OCRL. WDR81 has a peptide which does not fit in the consensus defined by the conservation of Ses1/2 F&H peptides (**Figure 1B**), but fits with the looser consensus (including an R residue at position 9) defined in **Figure 1C** by peptide array. This low-stringency F&H motif is lost in WDR81 in lower vertebrates such as *Gallus gallus* and *Danio rerio*, suggesting that it is not a conserved interactor of OCRL via this interface.

Our final candidate, Frabin/FGD4 is a Cdc42-GEF in the Dbl homology (DH) family of GEFs, consisting of a highly affine actin binding peptide region^26^, the DH-PH tandem GEF domain followed by two lipid binding domains: the PI3P-binding FYVE domain^27^ and a second PH domain, which binds a broad spectrum of phosphoinositides including PI(3)P and PI(4,5)P_2_^27^. As we had no antibody that recognised endogenous Frabin, we examined the interaction of Frabin-GFP with endogenous OCRL. Accordingly, a pool of Frabin-GFP was detected along with OCRL on spontaneously generated pinosomes in Cos7 cells (**Figure 1E**). It remained unclear whether this partial colocalization was due to a direct interaction or to the ability of both proteins to independently associate with pinosomes. However, when Cos7 cells expressing Frabin-GFP were subjected to anti-GFP immunoprecipitation, a weak, albeit specific, co-precipitation with OCRL was observed (**Figure 1F**). Modelling of Frabin using PHYRE2^28^ to thread the Frabin DH-PH sequence on the known structure of Vav (PDB:3BJI) suggests that the F&H peptide sits on an extended helix of the PH domain (**Figure 1G**), which may be concealed by the neighbouring FYVE domain when Frabin is inactive. The full-length Frabin-OCRL interaction may thus be regulated and only transient, despite the interaction of the Frabin F&H peptide with OCRL being strong.

Inspection of databases revealed that the F&H binding patch of OCRL is present in species in which APPL1, Ses1/2, and Frabin orthologues cannot be identified by primary amino acid similarity, suggesting that the original interactor of the OCRL-F&H interface had not yet been identified. Attempts to use unbiased bioinformatics approaches to identify conserved proteins harbouring F&H peptides failed, due largely to the large number of peptides that fit the putative F&H consensus and poor sequence conservation for domains such as PH or BAR domains. Thus, we turned to an ancient unicellular organism, *D. discoideum*, to isolate potential additional evolutionary conserved OCRL F&H interface interactors, and to understand the function of this interface.

### Identification of three F&H motif dependent interactors of Dd5P4

The single OCRL-like protein of *D. discoideum*, Dd5P4, has the same overall architecture as mammalian OCRL ^29,30^, except that it lacks the N-terminal PH domain^1^, and any identifiable clathrin coat binding motif (**Figure 2A** and **Supplementary Figure 2**). Importantly, the F&H motif binding surface of the RhoGAP-like domain is conserved. Dd5P4 knockout cells (*Dd5P4***^*-*^** mutants) can be partially rescued by expression of human OCRL ^29^, demonstrating that Dd5P4 and OCRL have well-conserved functions.

**Figure 2:**
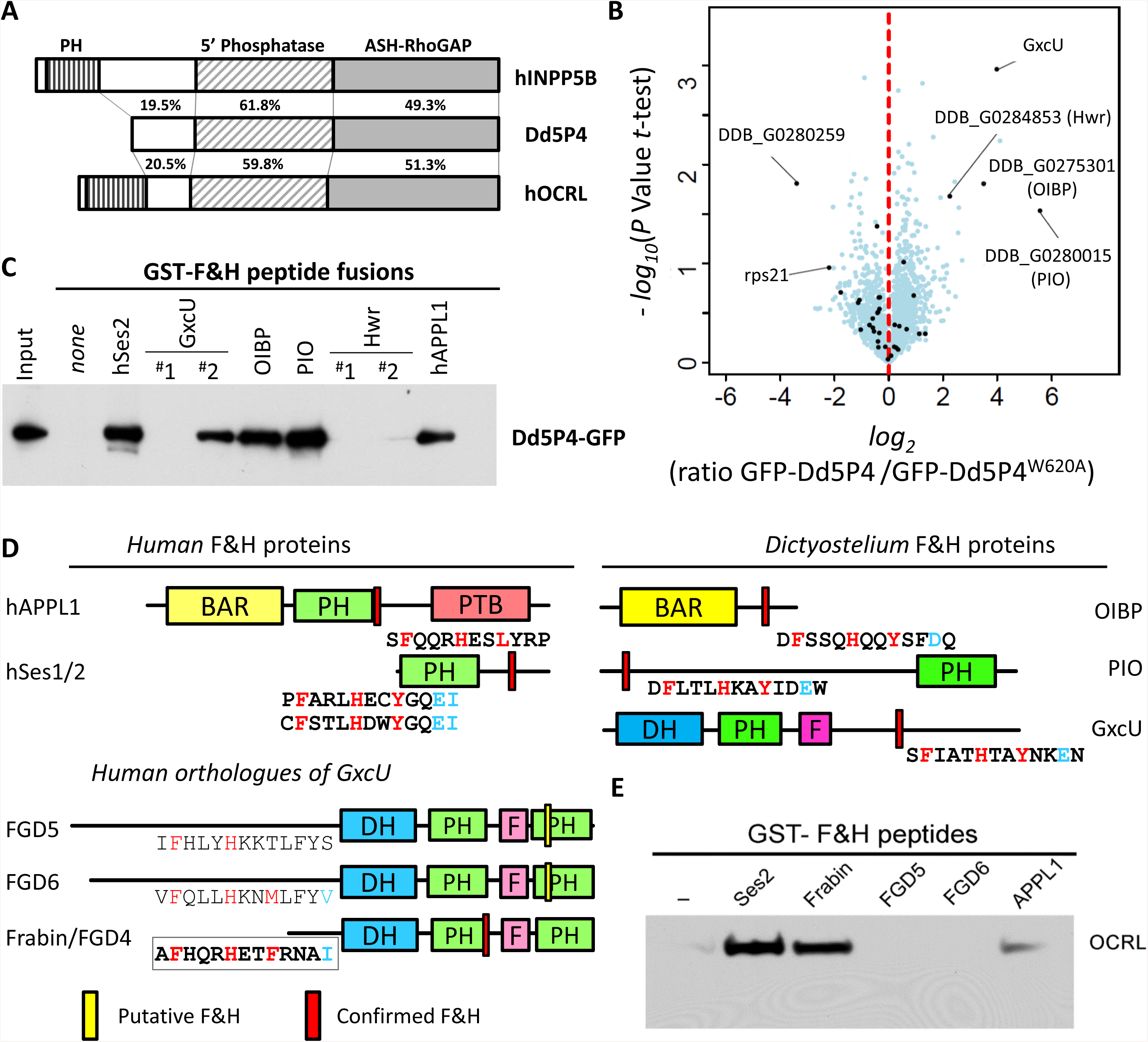
The F&H interface specifically interacts with three proteins in *D. discoideum*. **A)** Comparison of human INPP5B/OCRL and Dd5P4 amino acid sequences. **B**) Analysis of *Dd5P4*^*-*^; GFP-Dd5P4^WT^ vs *Dd5P4*^*-*^; GFP-Dd5P4^W620A^ immunoprecipitates shows enrichment of four possible interactors of the F&H surface of Dd5P4. Points labelled in Black are proteins that were also enriched in an analysis of wild-type vs *Dd5P4*^*-*^; GFP-Dd5P4^WT^ **(Supplementary Figure 3). C)** all possible F&H peptides in the four putative F&H-surface interactors were produced as GST fusions and purified on beads. These beads were exposed to lysates of *D. discoideum* expressing GFP-Dd5P4^WT^. **D)** Domain structure of F&H interactors in *D. discoideum* and humans, with validated interacting peptides shown below. *DH*: Dbl Homology domain, *PH*: Phox Homology homain, *F*: FYVE domain, *BAR*: Bin-Amphiphysin-Rvs domain, *PTB*: phosphotyrosine binding domain. The location of validated F&H peptide consensus is highlighted in red, unsuccessful F&H motifs are highlighted in yellow **E)** GST and GST-F&H peptides were exposed to HKC cell lysate. A specific interaction with endogenous OCRL was found with the known interactors Ses2 and APPL1.

To identify F&H interactors of Dd5P4 in *D. discoideum*, in *Dd5P4***^*-*^** cells we expressed GFP fusions of WT Dd5P4 and of a mutant Dd5P4 harbouring a tryptophan to alanine substitution at amino acid position 620, which corresponds to mammalian W739 required for F&H motif binding^4^ (**Supplemental Figure 2**). We then analysed triplicate GFP immunoprecipitates from untransfected wild-type cells, alongside GFP-Dd5P4 and GFP-Dd5P4^W620A^ transfected *Dd5P4***^*-*^**knockout cells by mass spectrometry. We found four proteins that were specifically enriched in precipitates from GFP-Dd5P4-expressing cells versus negative control precipitates but not in precipitates from GFP-Dd5P4^W620A^-expressing cells (**Figure 2B** and for comparison against GFP alone: **Supplementary Figure 3**). These four proteins were identified: three uncharacterized gene products, DDB_G0275301, DDB_G0280015, and DDB_G0284853 and the putative Rho-GEF GxcU.

To determine if there was a direct F&H peptide mediated interaction with Dd5P4-GFP, we made GST fusions of potential F&H peptides of each protein, performed pulldowns from lysates of *Dd5P4*^*-*^ cells expressing Dd5P4-GFP, and probed the blot with anti-GFP antibodies. To be certain that we captured any possible interaction, we selected as our test F&H peptide any 13aa stretch in the four proteins that contained the conserved F&H residues, including peptides which did not fit to either of our consensus motifs (**Figure 2C** and **Supplementary Figure 1**). GST-fusions of the F&H peptides of human Ses2 and APPL1 were also included. **Figure 2C** shows that Dd5P4-GFP bound the F&H motifs of human APPL1 and Ses2, confirming conservation of the F&H interaction between humans and *D. discoideum*. GST fusions of the putative F&H peptides of three of the four candidate *D. discoideum* interactors (DDB_G0275301, DDB_G0280015, and GxcU) bound Dd5P4 – while two low-stringency F&H peptides of protein DDB_G0284853, a mycBP2/Highwire-like protein (Hwr), did not (**Figure 2C, Supplementary Figure 1C**). We thus conclude that GxcU, DDB_G0275301 (which we called OCRL-**I**nteracting Bar domain Protein (OIBP)) and DDB_G0280015 (which we called PH domain **I**nteracting with **<u>O</u>**CRL (PIO)) - are *bona fide* direct interactors of the Dd5P4 F&H binding surface. Examining RNA expression profiles ^31^, all three F&H-containing proteins are well expressed in cells under non-differentiation conditions, suggesting that all three proteins are available to interact competitively with Dd5P4 at the same time.

### The three interactors of Dd5P4 have domain homologies to APPL1, Ses1/2 and Frabin

Remarkably, despite there being no identifiable amino acid sequence homology, modelling^28^ of the domain structures of the uncharacterized *D. discoideum* F&H motif interactors (OIBP, PIO and GxcU) suggested these to contain signature domains of those of the well-characterized F&H partners of OCRL: APPL1 (a BAR domain) and Ses1/2 (a PH domain with extended PxxP motifs), respectively. GxcU is a DH-PH domain Rho GEF with similar domain structure Frabin which we had identified as a human candidate interactor by peptide array (**Figure 2D**).

Using Phyre2^28^ to model *D. discoideum* sequences onto crystal structures deposited in the PDB database, OIBP (a.a 37-228) displayed a high structural homology with the BAR domain portion of the APPL1 paralogue APPL2 (PDB:4H8S, 98.8% confidence) in spite of a low (12%) sequence identity. The only published study referring to *D. discoideum* OIBP ^32^, reported that this protein was purified in association with macropinosomes, indicating a possible role in endocytosis/macropinocytosis, a function that is conserved with mammalian APPL1/2 ^10,18,33^.

PIO is a 741aa protein with a predicted PH-domain. Since PIO’s PH domain contains a series of long unstructured peptide loops that interfere with accurate tertiary structure prediction, to assess its structural “neighbours” we made structure predictions by searching only the portions of the PH domain of PIO conserved in the closely related species *Dictyostelium purpureum* (XP_003295168.1) (**Supplemental Figure 4**) ^34^. This PH domain falls loosely into the same structural family as the PH domain of PRKD3 (PDB:2D9Z 99.2% confidence 18% sequence similarity) and of PKCD2 (PDB:2COA; 99.2% confidence 17% sequence similarity), as does the PH domain of human Ses1 (99.9% model confidence, 17% sequence similarity). No function has previously been assigned to this protein.

GxcU is one of four *D. discoideum* proteins (GxcU, GxcV, GxcW and GxcX) with a domain structure similar to that of Frabin ^35^: a Dbl-Rho family GEF domain, followed by a PH domain and a FYVE domain. GxcU is largely unstudied, but knockout strains show that this gene is not essential for life^36^. Frabin (<u>F</u>GD1-related F-<u>a</u>ctin <u>bin</u>ding protein) is one of 7 closely related so-called FGD (faciogenital dysplasia) proteins, FGD1-6 and FRG (<u>F</u>GD1-related Cdc42-<u>G</u>EF)/FARP2, thought to function as Cdc42 GEFs. Three FGD proteins are associated with disease: FGD1 is mutated in Aarskog-Scott syndrome ^37^, FGD4/Frabin mutations cause the peripheral neuropathy Charcot Marie Tooth 4H^38^ and mutations of FGD6 are a risk factor for macular degeneration ^39^. Expanding on our peptide array analysis, we examined all putative F&H motifs in the FGD family regardless of possible presence in folded domains, isolating F&H peptides in Frabin/FGD4, FGD5 (a protein specific to haemopoetic stem cells ^40^) and FGD6 (**Figure 2D**).

Pulldowns from mouse brain extracts using GST fusions of the putative 13aa F&H motif peptides of each of the three FGD proteins, and of Ses2 and APPL1 as positive controls, revealed that the F&H peptide of Frabin, but not that of FGD5 and FGD6, specifically enriched endogenous OCRL (**Figure 2E)**. The F&H motif captured for Frabin was the same as that independently identified by peptide array (**Figure 1D**).

Two unbiased methods, peptide array on human proteins and mass spectrometry in *Dictyostelium* converge on a common set of conserved F&H-peptide harbouring OCRL interactors, APPL1, Ses1/2 and Frabin. We hypothesize that distant homologues of at least one of these proteins exist in all cells that express an OCRL protein harbouring this patch.

### F&H motif containing proteins are functionally similar in *D. discoideum* and mammals

We next examined the properties of our newly identified F&H binding partners in *D. discoideum*. PIO-GFP fusion proteins were located to compartments of the late-acidic/postlysosomal phase (**Figure 3**). PIO-GFP colocalised with the fluid-phase marker TRITC-dextran (a fluid-phase dye taken up in the phagosomal-lysosomal system and excreted via post-lysosomes) in organelles labelled by 30 minutes of fluid phase dye uptake by micropinocytosis, suggesting a function in the endo-lysosomal system. **Figure 3** shows PIO-GFP localization in both WT and *Dd5P4*^*-*^ cells exposed for 30 minutes to TRITC-Dextran. There is very little co-localization between PIO-GFP and dextran label at earlier timepoints (**Figure 3A**, ten minutes dextran pulse) and strong, but not complete colocalization at 30 minutes dye uptake (**Figure 3B –D)**, suggesting that PIO-GFP labels organelles at the transition between late acidic lysosomes to postlysosomes. This is reminiscent of the localization of Ses1/2 proteins at late endosomal stages in mammals ^10,19^.

**Figure 3:**
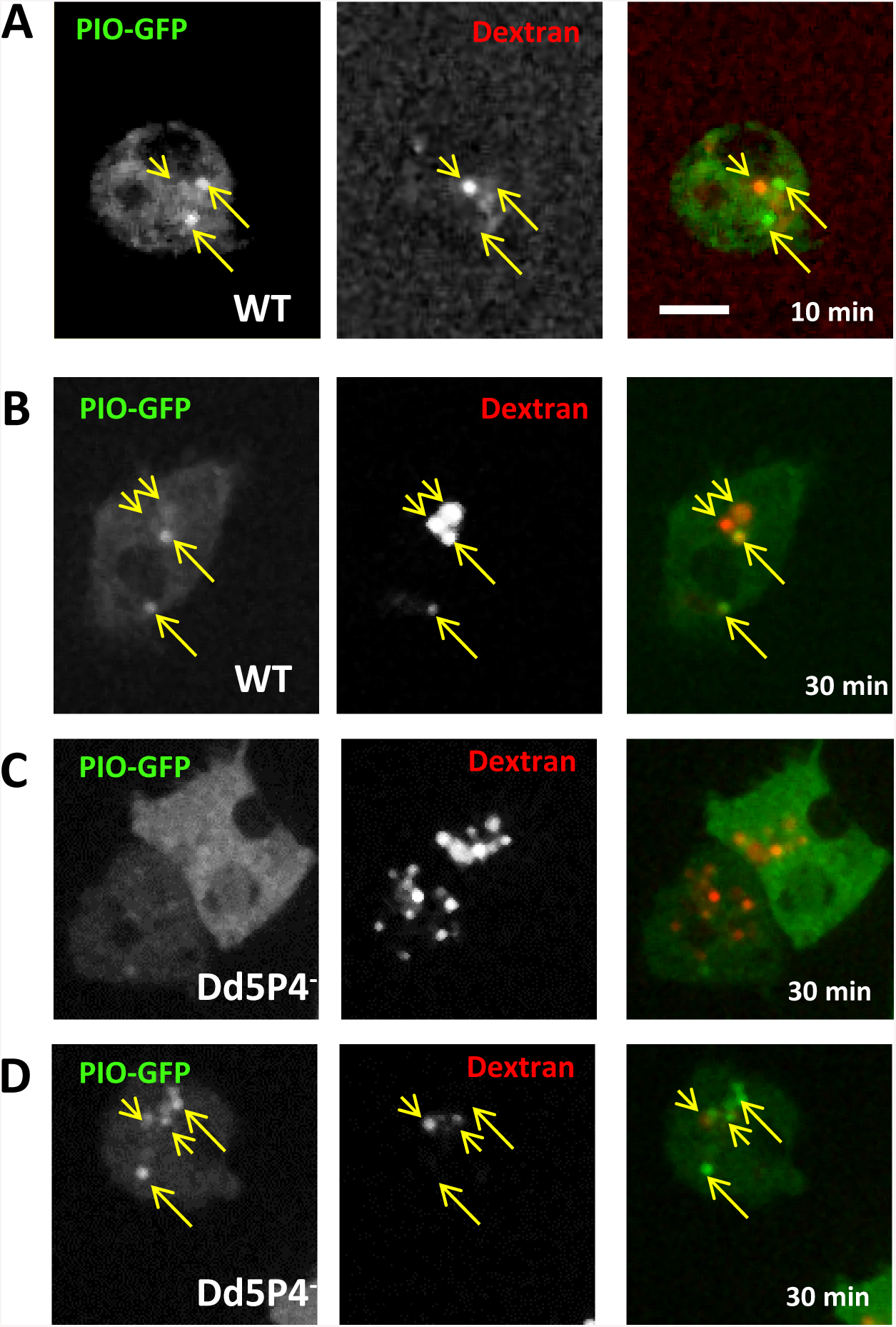
PIO-GFP labels a late lysosomal station. **A**) In WT cells PIO-GFP (*arrows*) does not colocalise with early lysosomal intermediates labelled by 10 minutes of fluid-phase dextran uptake (*arrowheads*) **B**) at 30 minutes, fluid phase dextran (*red*) marks some, but not all PIO-GFP positive compartments. **C**,**D**) There is no significant change in PIO-GFP localisation in *Dd5P4*^*-*^ cells: two examples of PIO-GFP expressing *Dd5P4*^*-*^ cells (one cytosolic, one punctate) labelled for 30 minutes with TRITC-dextran. Scale bar: 5μm.

Due to a long low complexity AT-rich stretch in the *D. discoideum* GxcU DNA sequence, it was not possible to clone or to synthetize the *D. discoideum* GxcU cDNA as a GFP fusion. Instead, we synthetized the sequence of the GxcU orthologue from the closely related amoeba *D. intermedium* ^34^, (see **Supplementary Figure 5** for sequence comparison) and expressed it as a fusion protein. Overall, GxcU-GFP appears more punctate in *Dd5P4*^*-*^ cells (**Figures 4A, B**). GxcU-GFP appears on several subcellular structures-including sites of macropinosome formation (**Figure 4C**) and on rare occasions at the plasma membrane some seconds after contractile vacuole (CV) exocytosis events (for a more complete description of CV, please see below) (**Figure 4D**), suggesting that GxcU participates in endocytic events. GxcU-GFP recruitment was evident at the base of the forming macropinocytic cup (**Figure 4C**), which would indicate an early endocytic function, and several seconds after CV fusion (**Figure 4C**) (*i.e*. later than the appearance of OIBP-GFP on these organelles (**Figure 5C**)). A role in coordinating actin-dependent remodelling of the plasma membrane at endocytic sites has also been reported for Frabin^41,42^. Recruitment of the GxcU-GFP fusion protein appeared delayed in the *Dd5P4*^*-*^ mutant (**Figure 4C**).

**Figure 4:**
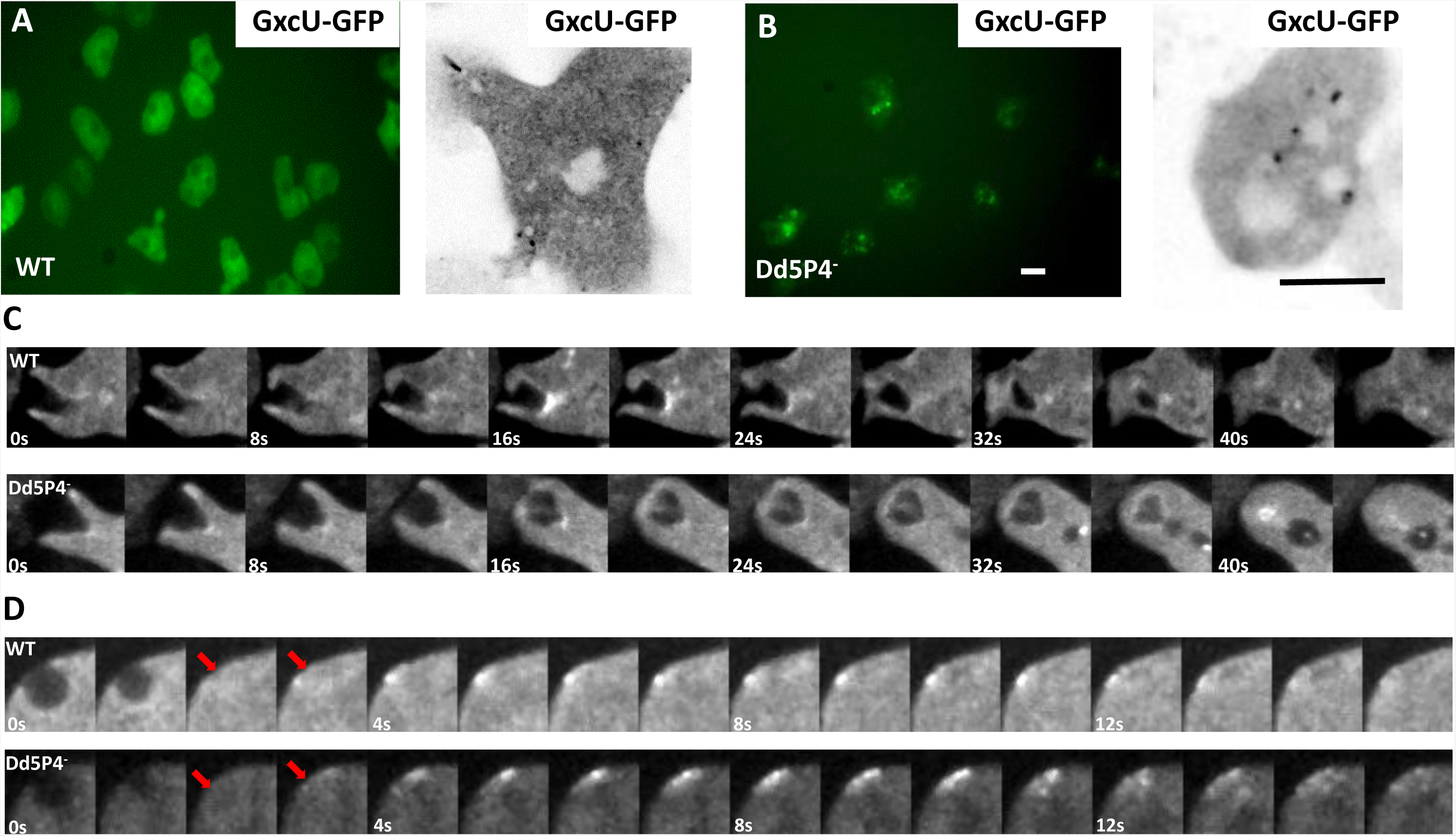
GxcU-GFP labels several endocytic intermediates. Wide-field microscopy (left panel) and confocal microscopy (right panel) of GxcU-GFP expressing cells **A)** wildtype cells expressing GFP-GxcU. **B)** GxcU**-**GFP is more punctate in *Dd5P4*^*-*^ cells. Scale bars in B: 10 µm. C) GxcU-GFP can be detected on phagosomes in both WT (upper) and *Dd5P4*^*-*^ (lower) cells. Gallery of images 0.25Hz (see **Supplementary Movies 2 and 3**) **D**) GxcU-GFP is also (rarely) detectible several seconds after CV collapse (site of CV collapse marked by red arrows) in both WT and *Dd5P4*^*-*^ cells, gallery at 1Hz.

**Figure 5:**
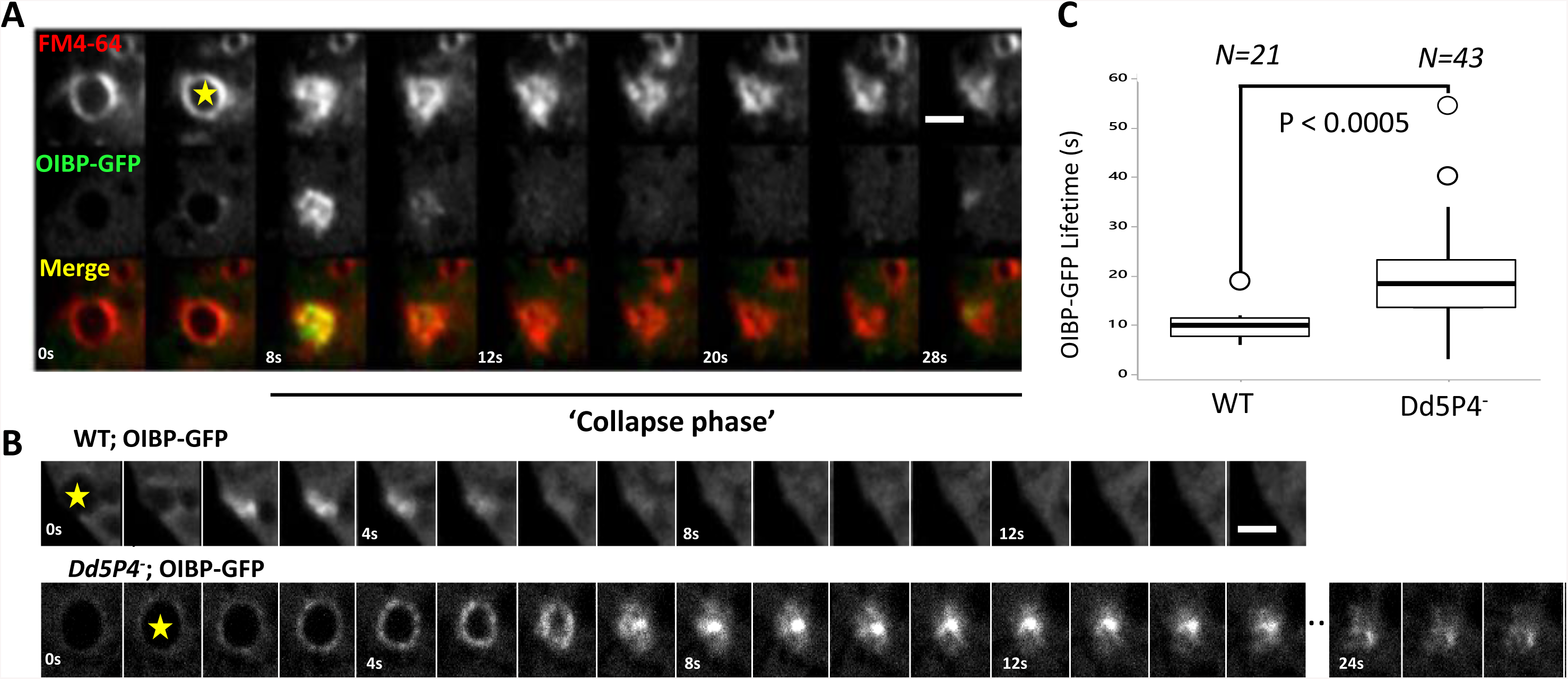
F&H interactions contribute to membrane reformation kinetics at the contractile vacuole. OIBP-GFP in **A**) wildtype cells. Expression of OIBP-GFP is largely cytosolic except for endocytic structures associated with contractile vacuoles, labelled in by FM4-64 (red). Shown is a gallery of images showing CV fusing *en face*, image rate 0.25Hz *Yellow Star*: CV beginning to collapse. (See **Supplementary Movie 4**). **B**) Gallery showing OIBP-GFP recruitment to osmotically-triggered CV fusion events imaged at 1Hz *upper panel:* WT cells. *Lower panel*: *Dd5P4*^*-*^ cells. Scale Bars in A and B: 4μm. **C**) Lifetime of OIBP-GFP fluorescence during CV collapse (start of the collapse indicated by yellow star) is approximately twice as long in *Dd5P4*^*-*^ cells. N=21 WT events, N=43 Dd5P4^-^ events, two sided student’s *T*-test: P<0.0005.

A C-terminal GFP fusion of OIBP was almost completely cytosolic except for brief and intense ‘flashes’ of recruitment, lasting only a few seconds, near the plasma membrane (**Figure 5A, B** and **Supplementary Movie 4**). As we could see no co-localisation of this signal with TRITC-dextran (*i.e*. the endolysosomal system), we examined another organelle, the contractile vacuole (CV). CVs act to excrete excess fluid to prevent osmotic rupture and are found in many water and soil dwellers. In *D. discoideum*, CVs are a reticulum of bladders and connecting tubules which fill with excess water, undergo rapid kiss-and-run fusion to disgorge their contents and locally recycle CV membrane without mixing with the plasma membrane by a striking process of tubulation, fragmentation and reformation of the CV organelle^43-45^ (here indicated as ‘collapse phase’). The lipophilic dye FM4-64 preferentially incorporates in CV membranes^46,47^. Co-labelling of the intensely OIBP-GFP-labelled structures with FM4-64 showed that these flashes occurred at the moment at which CVs open to the extracellular medium, during the discharge and collapse phase (**Figure 5A, B**). This observation suggests that OIBP-GFP is recruited as the CV membrane undergoes scission from the plasma membrane and is reformed by tubulation and fragmentation. This localization fits very well with the site of action in mammalian cells of APPL1, a protein with curvature inducer/sensor properties due its BAR domain^48^, which is recruited to endocytic membranes at the earliest stages of endocytosis in mammalian cells^18^. When this OIBP-GFP construct was expressed in *Dd5P4*^*-*^ cells, its localization was essentially unchanged (as is the case for its mammalian orthologue APPL1 in OCRL mutant patient fibroblasts^10^). However, when we examined the OIBP-GFP ‘flash’ at CV exocytic pores, we found that the lifetime of these events had approximately doubled in *Dd5P4*^*-*^ mutants (**Figure 5B** and **C**).

### Novel function of Dd5P4 in contractile vacuole fusion/endocytosis is modulated by F&H peptide interactions

Having determined that the F&H proteins of *D. discoideum* recapitulate the localization of their mammalian counterparts, we turned to characterize Dd5P4 recruitment and function on these organelles. Complementation of a *Dd5P4*^*-*^ line with Dd5P4 fused to GFP at either the N- or C-terminus rescued previously-characterized growth deficits^34^ in these mutants, indicating that the fusion proteins were functional (**Supplementary Figure 5A**). We also successfully rescued the *Dd5P4*^*-*^ growth deficit with the F&H interface mutant GFP-Dd5P4^W620A^, suggesting that either sufficient overexpression of Dd5P4^W620A^ is enough to overcome any defects in efficient targeting of Dd5P4 activity to membranes, or that, as we had previously observed in human patient fibroblasts^4^, loss of the F&H interface led to loss of OCRL-like activity from very specific membrane subcompartments, and not a generalised deficit in OCRL-mediated traffic.

Given the presence of GxcU-GFP and PIO-GFP in the micropinocytic system and the functional requirement for Dd5P4 for micropinocytosis/nutrient uptake (**Supplemental Figure 5A**), we re-examined the subcellular recruitment^29^ of fluorescently tagged Dd5P4. Despite an established function of Dd5P4 in phagocytosis^29^, and despite the recruitment of mammalian OCRL to endocytic vesicles engulfing pathogens^49^, neither published studies ^29^ nor our own study with N or C-terminal GFP-tagged Dd5P4 fusions detected a clear accumulation of Dd5P4 on macropinocytic intermediates (labelled by TRITC-dextran in **Supplementary Figure 5B**). We hypothesise that GFP-tagged fusion proteins are likely to be expressed at higher levels than endogenous Dd5P4, and the excess cytosolic fluorescence may mask the presence of a macropinosome-associated pool of the protein.

*Dd5P4*^*-*^ cells expressing either GFP-Dd5P4^WT^ or GFP-Dd5P4^W620A^ exhibited marked transient recruitment of GFP fluorescence to organelles that appeared to fuse with the plasma membrane. We excluded post-lysosomes^50,51^ as the organelles in question, as GFP-Dd5P4^WT^ and GFP-Dd5P4^W620A^ fluorescence segregated away from post-lysosomal organelles when cells had been preloaded overnight with TRITC-dextran (**Supplementary Figure 5B**). FM4-64 dye labelling showed both GFP-Dd5P4^WT^ and GFP-Dd5P4^W620A^ were recruited to CVs in the moments before its fusion and collapse at the plasma membrane (**Figure 6D**). We then explored the functional requirement for Dd5P4 and its F&H interface on this organelle.

**Figure 6:**
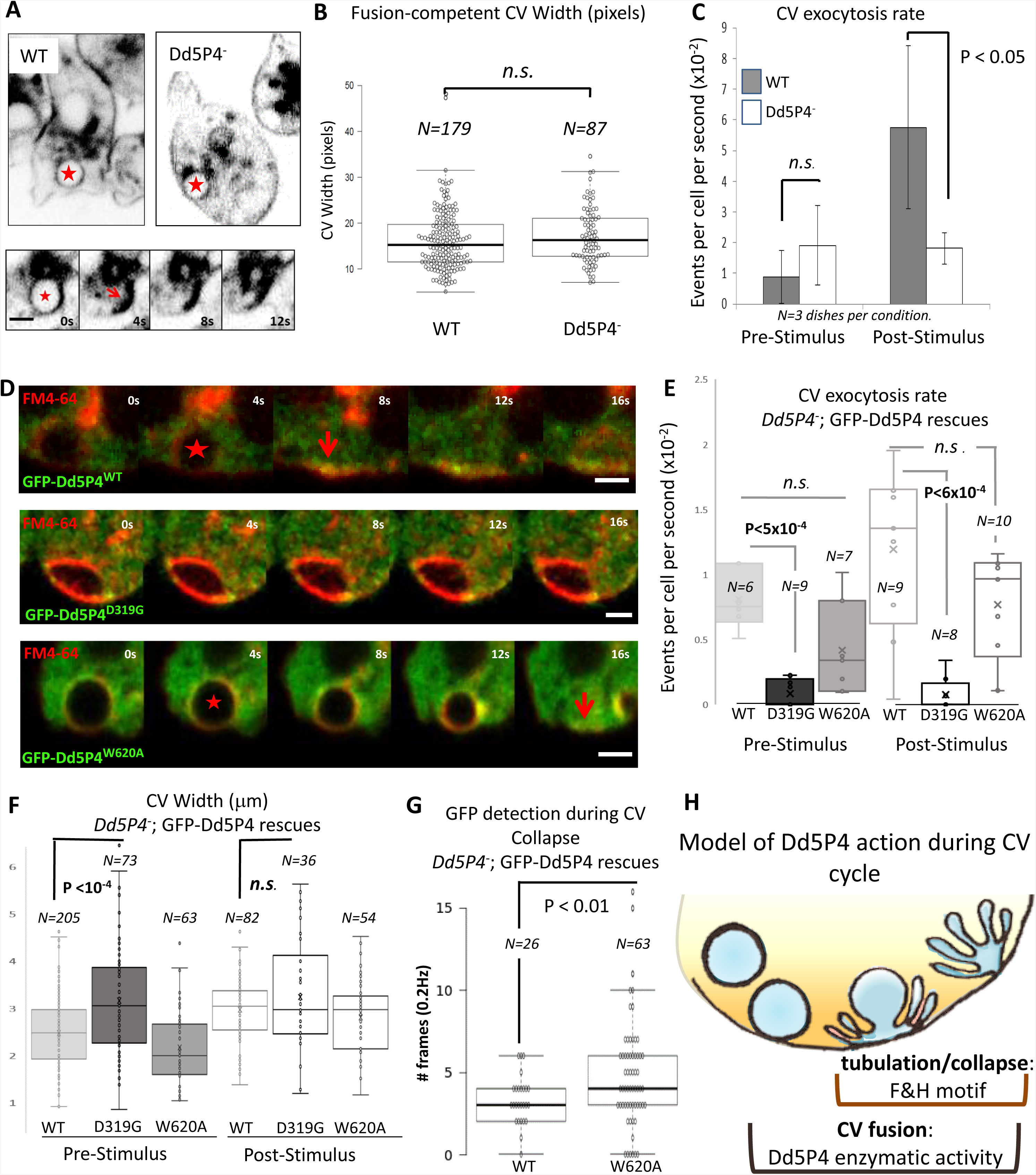
CV dynamics in Dd5P4 mutants and rescues **A)** FM4-64 dye labelling of CV membranes in WT and *Dd5P4*^*-*^ cells show CV distribution is similar. Lower panel: CV collapse event (frames from 0.25 Hz movie), quantified in **6C** and **E**. *Star* indicates a CV about to collapse, *arrow* indicates the collapsed CV membrane. **Scale bar:** 2μm **B)**. Size at widest point of CVs undergoing fusion is unchanged by Dd5P4 loss: N=179 (WT), =87 (Dd5P4^-^). **C)**. rate of CV fusion triggered by 1:1 osmotic stimulus is strongly reduced, but not abolished, in *Dd5P4*^*-*^ mutants. Average of 3 movies per condition, StdDev, two-sided student’s *T*-test.**D**) Recruitment of GFP-Dd5P4 constructs to contractile vacuoles marked by FM4-64. GFP-Dd5P4 ^WT^ and GFP-Dd5P4^W620A^ are recruited to CVs at the moment that the CV starts to collapse (marked by a *red star*) and persists at the moment of full collapse (*arrow*). GFP-Dd5P4^D319G^ is recruited in punctae to the CV but do not appear to spread over the CV surface, CVs appear enlarged. Movie: 0.25Hz**. Scale bars:** 2μm**. E)** FM4-46 labelling shows that *Dd5P4*^*-*^; GFP-Dd5P4^D319G^ cells exhibit significantly reduced CV fusion/collapse in both basal and osmotically stimulated conditions. *Dd5P4*^*-*^; GFP-Dd5P4^W620A^ cells show a similar rate of CV fusions as *Dd5P4*^*-*^; GFP-Dd5P4^WT^. number of movies per condition indicated on graph. Student *T*-Test. See **Supplementary Movies 7 and 8** for live imaging of GFP signal. **F)** Cells in basal conditions show significantly larger CV diameters in *Dd5P4*^*-*^; GFP-Dd5P4^D319G^ cells. **G)** GFP signal in collapse phase lasts longer in *Dd5P4*^*-*^; GFP-Dd5P4^W620A^ cells than in cells re-expressing GFP-Dd5P4^WT^. **H**) Model of Dd5P4 action during CV cycle

When we imaged FM4-64-labelled cells, we identified frequent CV discharges (see **Figure 6A** for examples of fusion events, showing a full CV, collapse of the organelle and local membrane tubulation). The size of CVs that underwent exocytosis was unchanged between wildtype and *Dd5P4*^*-*^ cells (**Figure 6B**), suggesting that CV biogenesis was grossly normal. Additionally, basal CV fusion, unstimulated by osmotic pressure, occurred at the same rate in both WT and *Dd5P4*^*-*^ mutant strains. However, 1:1 dilution of the imaging medium with distilled water caused an increase in water pumping and of the CV exocytic rate in WT (**Figure 6C Supplementary Movie 5**), whereas the exocytic rate of CVs did not change from the basal rate in the *Dd5P4*^*-*^ mutants (**Supplementary Movie 6**).

Re-expression of GFP-Dd5P4^WT^ in *Dd5P4*^*-*^ mutants rescued the osmotically-triggered component of CV exocytosis observed in these mutants (**Supplementary Figure 5C**). GFP-Dd5P4^W620A^ showed localization on CVs similar to GFP-Dd5P4^WT^, speaking against a role of F&H-containing proteins in the recruitment of Dd5P4 to the CV membrane. Comparing the re-expression of GFP-Dd5P4^WT^ in *Dd5P4*^*-*^ mutants with that of GFP-Dd5P4^W620A^ or the catalytic mutant GFP-Dd5P4^D319G^, (**Figure 6D**) we noted that the catalytic mutant exerted a dominant effect, almost completely abolishing CV cycling (**Figure 6E**) in both resting and osmotically-stimulated conditions. Additionally, we observed that CV diameter (FM4-64 staining) increased in resting cells expressing GFP-Dd5P4^D319G^ (**Figure 6D**, quantified in **Figure 6F**). Interestingly, whereas wildtype and W620A Dd5P4 appeared to be recruited to the entire CV membrane at about the time when CV starts decreasing in size due to CV discharge (stars in **Figure 6D**) and persists during the tubulation/collapse phase (arrowheads in **Figure 6D**), GFP-Dd5P4^D319G^ appeared punctate, and did not spread across the CV membrane upon arrival at a FM4-64-labelled organelle. We did not capture any events where GFP-Dd5P4^D319G^ was recruited to a collapsed/tubulating CV. We therefore analysed only the kinetics of GFP recruitment in *Dd5P4*^*-*^;GFP-Dd5P4^WT^ or *Dd5P4*^*-*^;GFP-Dd5P4^W620A^ CV fusion events. There was no significant difference in the lifetime of the GFP signal on pre-collapse CVs between the two fusion proteins (**Supplemental Figure 5D**), as in general both fluorescent proteins were recruited when the CV started to discharge and collapse (a few CVs took some time to collapse after GFP-Dd5P4 recruitment as shown in **Supplementary Figure 5D**). We found that GFP-Dd5P4^W620A^ lingered for a significantly longer time than GFP-Dd5P4^WT^ during the process of CV collapse/tubulation (**Figure 6G**) suggesting defects in the process by which membranes tubulate and reform after CVs discharge their content. Taken together, this indicates that catalytic activity is necessary for Dd5P4-mediated CV fusion and that F&H peptide interactions in *D. discoideum* are not essential for recruitment of Dd5P4 to CV membranes, but are necessary for efficient dynamics/remodelling of these membranes during fusion and subsequent re-endocytosis of CV membranes (model in **Figure 6H**).

### Dd5P4 loss-of-function deficit in fluid phase dye uptake is phenocopied and enhanced by GxcU overexpression

Loss of Dd5P4 has been shown to cause deficits in fluid-phase micropinocytosis. Neither PIO-GFP nor OIBP-GFP appeared to be recruited to early micropinocytic intermediates, but GxcU-GFP had a wide distribution on organelles including micropinosomes (**Figure 4**), suggesting the two proteins may interact in this process. Overexpression of GxcU-GFP inhibited uptake of TRITC-dextran (**Figure 7A, B**), over all the timescales measured (10minutes, 60 minutes and overnight application of labelled dextran to the culture medium), as observed with Dd5P4^-^ mutants. Both Dd5P4^-^ mutants and GFP-GxcU overexpressors exhibited numerous small dye-positive endolysosomal structures rather than the large lysosomes/postlysosomes found in WT cells, suggestive of a defect in membrane traffic along multiple parts of the endo/lysosomal pathway (see inset images at 60 mins uptake in **Figure 7A**). GxcU-GFP overexpression in Dd5P4^-^ evoked a profound deficit in dye uptake, suggesting that Dd5P4 activity opposes the action of overexpressed GxcU (**Figure 7B, C**). Cumulatively, this data suggests that OCRL and its F&H protein interactors each act defined points of the membrane cycle as illustrated in **Figure 7D**.

**Figure 7:**
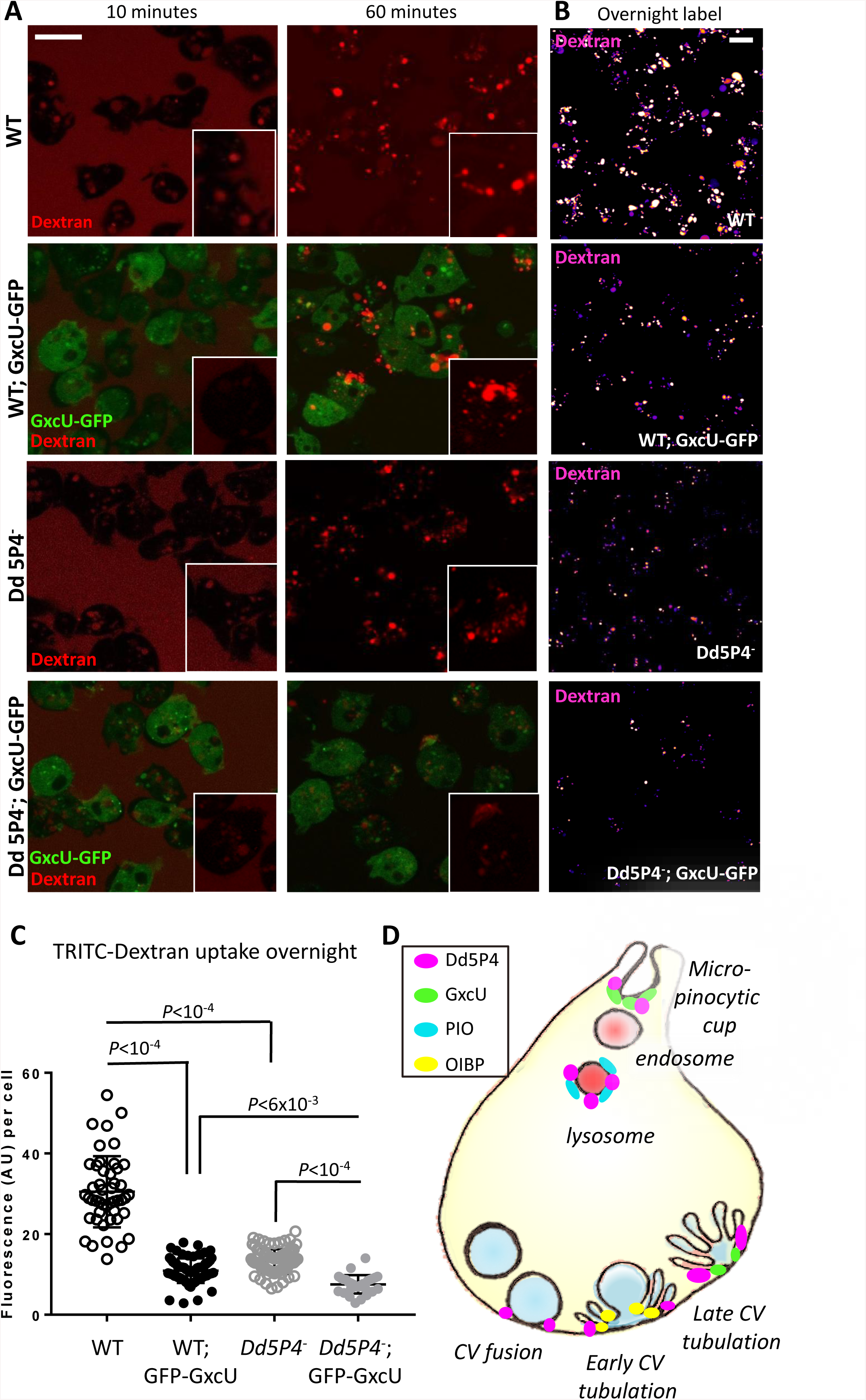
GxcU-GFP overexpression and Dd5P4 loss show similar effects in micropinocytic dye uptake and subsequent membrane trafficking. **A**) timed application of TRITC-dextran (red) shows defects in initial micropinocytic uptake (10 minutes) and subsequent membrane trafficking (60 minutes) in both *Dd5P4*^*-*^ cells and GxcU-GFP overexpressors, which is enhanced by GxcU-GFP overexpression in the *Dd5P4*^*-*^ background. Individual cells are shown *inset*. **B**) Deficits in dye uptake are not resolved by longer exposure to dye (overnight labelling). TRITC-dextran label is shown in false colour scale for ease of comparison. Scale Bars in **A, B**: 8μm. **C**) GFP-GxcU overexpression enhances dye uptake defects caused by Dd5P4 loss. Fluorescence per cell after overnight TRITC-dextran uptake, ANOVA.**D**) localisation of Dd5P4 and F&H-motif-containing interactors

## Discussion

Previous studies have shown that interactions of F&H-peptide-containing proteins with OCRL/INPP5B participate in the localization and function of these inositol 5-phosphatases on intracellular membranes^4,19^. Loss of the F&H-binding surface in mammals very subtly re-localizes OCRL, whereas loss of the OCRL Rab binding interface^4,52^ almost completely delocalizes the protein, suggesting that the F&H interface, as we see in *D. discoideum*, has a specialized function in membrane traffic, rather than a generalized function recruiting OCRL. The F&H motif containing OCRL/INPP5B interactors APPL1 and Ses1/2 (IPIP27A/B), were thought to appear later in evolution than OCRL, as searches of databases had detected APPL1 only in vertebrates^3^ and Ses/CG12393 in *Drosophila* and other insects, but not in unicellular organisms ^10^. In contrast, OCRL/INPP5B-like proteins are widely present in unicellular organisms. Amino acids critically required for the F&H interacting surface of OCRL^4^ were highly conserved even in these ancestral OCRL species, suggestive of the presence of yet-to-be-identified F&H interactors in these evolutionary older organisms.

We tested this possibility in the social amoeba *D. discoideum*, which expresses a single OCRL orthologue, Dd5P4, displaying an F&H interacting surface. Loss of Dd5P4 is not lethal ^30^, but leads to quantifiable defects in macropinocytosis and growth ^29^, which can be rescued by expression of GFP-fusions of Dd5P4 and of human OCRL, suggesting closely conserved function and subcellular targeting^29^.

To our surprise, we found that the identity of the F&H proteins that interact with OCRL is as well-conserved as the F&H surface itself: in both humans and amoeba, OCRL uses the F&H interface to couple to three proteins, a BAR-domain protein present at the earliest stages of endocytosis (APPL1 in humans and OIBP in *D. discoideum*), a PH-domain containing protein on late lysosomes/postlysosomes (Ses1/2 in humans and PIO in *D. discoideum*), and a novel interactor GxcU/ Frabin, a Rho family GEF, which in humans is associated with the disorder Charcot Marie Tooth Type 4. Live imaging of GxcU and Frabin showed that both proteins appear at multiple early and later stages of endocytosis. Our data suggests that there is a specific requirement of OCRL phosphatase activity coordinated by different F&H proteins at well-delineated endocytic stations.

We also established a novel function for OCRL-like phosphatases in membrane trafficking: in addition to known phenotypes of Dd5P4 loss in *D. discoideum* ^29,30,49,53^, (reduced growth and a defect in both micropinocytosis and phagocytosis), we identified a novel phenotype in axenic cells: Dd5P4 controls the rate of fusion and endocytic tubulation/recapture of an osmoregulatory organelle, the contractile vacuole (CV). CV exocytosis is thought to be a “kiss-and-run” process, involving minimal mixing of the CV membrane with the plasma membrane^54^.

Notably, we found that Dd5P4 has two separable functions in the CV discharge process: CV fusion, which did not depend on F&H peptide interactions but was blocked by overexpression of a catalytically inactive Dd5P4 fusion protein, and tubulation/reformation (which we indicate as ‘collapse phase’), whose kinetic was influenced by F&H-mediated interactions. We measured extended kinetics of the CV ‘collapse phase’ in GFP-Dd5P4^W602A^ expressing mutants (where the CV fusion rate defects in Dd5P4 knockouts is restored), and the extended kinetics of the OIBP-GFP fusion protein in this same stage in *Dd5P4*^*-*^.

How Dd5P4 interferes with CV fusion is yet unclear, although several functions of OCRL/Dd5P4 might be implicated. Disturbances of the actin cytoskeleton cause inappropriate mixing of CV and plasma membranes ^45^, leading to CV-resident proton pumps being found at the plasma membrane. Similarly, ‘spillover’ of the transmembrane protein Dajumin, responsible for the biogenesis of CVs, is retrieved from the plasma membrane *via* clathrin/AP2^55^, where mammalian OCRL is also known to act. Thus it is possible that Dd5P4 could have several functions on this organelle, for example regulating underlying actin and/or PI(4,5)P_2_ at the CV fusion site, or in trafficking of receptors which are necessary for the CV fusion/reformation cycle.Our images of GFP-Dd5P4^D419G^ show that this mutant protein remains clustered in punctae upon arriving to docked CVs, whereas GFP-Dd5P4^WT^ and GFP-Dd5P4^W620A^ arrive at CVs prior to fusion and then spread over the CV membrane, coincident with the CV starting to fuse and decrease in size. Previous work ^54,56^ suggests that during this phase of the CV cycle (docking and fusion-ready) CVs are labelled by Drainin and Rab8 and undergo ‘ring to patch’ transition to allow full fusion. In the absence of the ‘ring to patch’ transition (f.e. in P2XA mutant CVs^56^) CV fusion becomes inefficient, as we have seen in *Dd5P4*^*-*^ mutants. In contrast, the more severe phenotype of GFP-Dd5P4^D419G^ suggests that this mutant may sequester factors such as Rab8^GTP^. Indeed the swollen CV phenotype found in GFP-Dd5P4^D419G^ is reminiscent of the Rab8^DN^ and *drainin*^*-*^ phenotypes previously published^54^.

While there is no direct analogue of this organelle in mammals, the CV is decorated by the small GTPase Rab8 prior to its exocyst-directed fusion with the plasma membrane ^54^. Rab8 is a strong interactor of OCRL in mammals ^57,58^, where the interaction is responsible for membrane traffic to the primary cilium^21^. Interestingly, both CV fusion in *D. discoideum* ^54^ and growth of cilia involve fusion *via* the exocyst complex ^59^, suggesting OCRL may serve an analogous function in both contexts.

The PH domain protein PIO, like its higher-organism counterpart Ses1/2, is a late endocytic protein, only being present on endocytic organelles that are reached by fluid-phase endocytic tracer 30 minutes or more after ingestion (**Figure 3**), consistent with a late-endosomal or lysosomal function. The localization of the newly identified OCRL/Dd5P4 interactor, GxcU/Frabin, was particularly intriguing. In human cells, Frabin-GFP has a similar multi-stage recruitment pattern to both early and late endosomal and macropinocytic membranes^60,61,62^ (**Figure 2, Supplementary Movie 1**). In *D. discoideum*, GxcU-GFP was recruited to several distinct stages of the endocytic pathway (**Figures 4E and F**), both at what would be considered very early (at the base of the macropinocytic crown) and at the site of reforming CVs, several seconds after the timepoint when the other F&H interactor, OIBP-GFP appears (*e.g.* **Figure 5A**). While the target of the GxcU GEF activity is not known, the localization of GxcU to macropinosomes is highly reminiscent of the localization of active Rac1 ^63,64^. Interestingly, overexpression of GxcU-GFP phenocopied Dd5P4 loss in terms of fluid-phase dye uptake and processing and enhanced the Dd5P4^-^ mutant phenotype (**Figure 7**), suggesting that Dd5P4 may be a repressor of GxcU function.

In summary, our findings reveal that the network of F&H peptide interactors of OCRL is more evolutionary conserved than expected based on previous studies. Furthermore, in all cases that we examined, selective disruption of the F&H interface of OCRL/Dd5P4 does not significantly impair its localization, but impacts downstream reactions. In *D. discoideum*, F&H mutant Dd5P4 localizes to the same subcompartments as wildtype Dd5P4, but where we made kinetic measurements, we found that lack of F&H interactions affected the kinetics of the trafficking step under study. Likewise F&H-containing interactors of Dd5P4 were recruited to membranes for longer periods in *Dd5P4*^*-*^ mutants than in wild type, suggesting that co-ordinated activity of OCRL and F&H adaptors is necessary for efficient membrane progression.

Our study gives weight to the possibility of functionally relevant amounts of PI(4,5)P_2_ being present in the endocytic system under normal conditions. Recent research suggests that PI(4,5)P_2_ and/or OCRL catalytic activity has a critical role in membrane traffic separate from its role in at the plasma membrane, including transition from early to late endosomes^65-68^ and the transition from endosome to trans-Golgi^8,22^. The presence of conserved, endomembrane-specific interactors for OCRL argues against a function of OCRL solely as a cellular ‘housekeeper’, and suggests instead that OCRL-directed PI(4,5)P_2_ on endosomes is assisted and directed by interaction with F&H proteins.

Fibroblasts from OCRL- mutated Lowe Syndrome or Dent2 patients are equally deficient in PI(4,5)P_2_ phosphatase activity ^69^, however Dent2 disease fibroblasts manifest milder phenotypes (*e.g.* the degree of defective actin stress fibre formation, abnormal alpha-actinin staining and shortened primary cilia^69^) than their Lowe syndrome counterparts, suggesting the presence of a modifier gene on an autosome. The OCRL paralog INPP5B has been excluded as a phenotype modifier in Lowe/Dent2 patients ^69^. Given that the F&H interaction is so well conserved and appears to change the kinetics of OCRL-dependant membrane trafficking steps, F&H proteins should be considered as highly likely candidates for modifiers of the Lowe syndrome phenotype.

## Acknowledgements

LES was supported for this project by Wellcome Trust (105616/Z/14/Z), Medical Research Council (MRC/N010035/1) and Lowe Syndrome Association USA. PDC was supported by NIH grants (DA018343; DK082700; NS036251), the Lowe Syndrome Trust, the Lowe Syndrome Association USA and the Kavli Institute for Neuroscience. TS laboratory is supported by multiple grants from the Swiss National Science Foundation. TS is a member of iGE3 (www.ige3.unige.ch). GC was supported by the Italian Association for Cancer Research Grant N (IG 2013 N.14135). MS is supported by North West Cancer Research (CR1081).

## Author Contributions

AL, FF, CB, CL, JM,FL, AP, MP, LES performed experiments. AL FF CB FL JM, AP MP MS GC TS, PDC and LES analysed data, FF CB TCW TS PDC and LES contributed to the writing of the manuscript.

---

**Supplementary Figure 1, related to Figure 1.** Determination of candidate F&H proteins in humans by peptide array. **A**) Western Blot on three candidate proteins in HKC lysates showed no specific interaction with the OCRL F&H interface. **B**) ITC reveals the putative F&H peptide of Dynein heavy chain to be a non-specific interactor. **C**) Summary of tested human F&H peptides. Proteins labelled in red have been validated in previous publications. Proteins in yellow have been confirmed to be non-interactors of the OCRL-F&H motif by Isothermal Titration Calorimetry. Red residues are common to Ses1/2 and APPL1, blue residues are conserved in Ses proteins. Purple residues are at key positions and fall outside the Ses/APPL1 consensus. (MS= mass spectrometry on GFP-OCRL precipitates), WB (Western Blot) ITC (Isothermal titration calorimetry). + published in Swan et al PNAS 2010 and Pirruccello et al NSMB 2011; * Nandez et al eLIFE 2014.

**Supplementary Figure 2, related to Figure 2**. Sequence conservation between Dd5P4 and human INPP5B and OCRL. Domains are indicated by lines under the sequence alignment. *Identical residues* are shaded. *Boxed residues*: interaction motifs for clathrin and AP2 (unique to OCRL). *Dot*: catalytic residue. *Arrowhead:* key Rab binding residue. *Starred residues*: mediate contact with F&H peptides. *Residues marked in red:* mutated in this study.

**Supplementary Figure 3, related to Figure 2**: Analysis of peptide enrichment of GFP immunoprecipitates from **A**) WT vs *Dd5P4*^*-*^;GFP-Dd5P4^WT^ and **B**) WT vs *Dd5P4*^*-*^;GFP-Dd5P4^W620A^ shows that putative F&H proteins (**Figure 2B**) are specifically enriched in precipitates from GFP-Dd5P4^WT^ expressing cells.

**Supplementary Figure 4**: related to **Figures 3 and 4**. Sequences of PIO and GxcU proteins **A)** The predicted PH domains of *D discoideum* PIO (aa 543-741) and *D. purpureum* PIO partial sequence (aa 200-401). Identical residues are shaded. *Boxed region* was used to search crystal structures with Phyre2 **B)** Conservation of amino acid sequence between *D discoideum* and *D intermedium* GxcU genes. The Rho-GEF, PH and FYVE domains are strongly conserved, as is the F&H peptide sequence (boxed residues).

**Supplementary Figure 5, related to Figure 6**: GFP-Dd5P4^W620A^ rescues many aspects of *Dd5P4*^*-*^ dysfunction. **A**) N-terminal and C-terminal GFP fusions of Dd5P4, and an N-terminal GFP labelled mutant of the F&H surface (W620A) all restore growth in a *Dd5P4*^*-*^ background. N=3 independent growth curves per genotype. **B**) Both GFP-Dd5P4 and GFP-Dd5P4^W620A^ are largely cytosolic (as seen in previous publications), but GFP fluorescence is found on cytosolic organelles which are not part of the endocytic system (negative for 7KDa Dextran label).Arrowheads show GFP accumulation at collapsing contractile vacuole. **C**) Re-expression of GFP-tagged Dd5P4 restores osmotically-triggered CV fusion in cells labelled with FM4-64, N=3 independent movies per genotype, normalised to corresponding wildtype value. **D**) Lifetime of GFP-labelled CVs imaged at 0.2Hz *prior* to fusion in *Dd5P4*^*-*^; GFP-Dd5P4^WT^ cells is not significantly different to that of *Dd5P4*^*-*^; GFP-Dd5P4^W620A^ expressers: n= 28 GFP-Dd5P4^WT^ labelled events, n=67 GFP-Dd5P4^W620A^ labelled events, Student’s t-test. All CV exocytosis events which were GFP positive either in the pre-collapse or collapse phase were analysed. The majority of GFP-Dd5P4^WT^ CVs are positive for GFP only in the collapse phase (i.e are recorded as zero frames before CV collapse).

**Table 1:** constructs and *Dictyostelium* cell lines used in this manuscript

**Supplementary Movie 1, related to Figure 1:** Ruby-OCRL (red) and Frabin-GFP (green) are found together on pinosomal compartments in Cos7. Images 0.25Hz. Scale Bar 3μm.

**Supplementary Movie 2, related to Figure 4:** GxcU-GFP can be detected on phagosomes in WT cells. Images 0.25Hz. Scale Bar 15μm.

**Supplementary Movie 3, related to Figure 4:** GxcU-GFP can be detected on phagosomes in *Dd5P4*^*-*^cells. Images 0.25Hz. Scale Bar 15μm.

**Supplementary Movie 4, related to Figure 5:** Recruitment of a C-terminal GFP fusion of OIBP (green). Contractile vacuoles are labelled by FM4-64 (red). Images 0.25Hz. Scale Bar 10μm.

**Supplementary Movie 5, related to Figure 6**: 1:1 dilution of imaging medium with distilled water caused an increase CV exocytosis rate in WT. Images 0.2 Hz. Scale Bar 15μm.

**Supplementary Movie 6, related to Figure 6:** 1:1 dilution of imaging medium with distilled water in *Dd5P4*^*-*^ mutants does not change CV exocytosis rate. Images 0.2 Hz. Scale Bar 15μm.

**Supplementary Movie 7, related to Figure 6:** the collapse/fusion phase of CV exocytosis in *Dd5P4*^*-*^ mutants re-expressing GFP-Dd5P4. Images 0.2Hz. Scale Bar 15μm.

**Supplementary Movie 8, related to Figure 6:** the collapse/fusion phase of CV exocytosis in *Dd5P4*^*-*^ mutants re-expressing F&H mutant GFP-Dd5P4^W620A^. Images 0.2Hz. Scale Bar 15μm.

