## Supplementary material for "Lowe Syndrome-linked endocytic adaptors direct membrane cycling kinetics with OCRL in *Dictyostelium discoideum*": Table 1

**Table 1. *Dictyostelium* material used within this publication.**

| ***Dictyostelium* strains used** | | **Plasmids used for transformation** | **Reference** | |
| --- | --- | --- | --- | --- |
| AX3 (wild type) | |  |  | |
| *dd5P4* knockout strain | |  | (Loovers et al., 2007; Loovers et al., 2003) | |
| GFP-Dd5P4/*dd5P4* KO | | see below |  | |
| Dd5P4-GFP/*dd5P4* KO | | see below |  | |
| GFP-Dd5P4^W620A^/*dd5P4* KO | | see below |  | |
| GFP-Dd5P4^D319G^/*dd5P4* KO | | see below |  | |
| OIBP-GFP/ *dd5P4* KO | | see below |  | |
| PIO-GFP/ *dd5P4* KO | | see below |  | |
| GxcU-GFP/ *dd5P4* KO | | see below |  | |
| **Plasmids generated for this work** | **Insert** | | | **Plasmid &Reference** |
| Dd5P4-GFP | DDB_G0267462 | | | pDM323 (Veltman et al., 2009) |
| GFP-Dd5P4 | DDB_G0267462 | | | pDM317 (Veltman et al., 2009) |
| GFP-Dd5P4^W620A^ | DDB_G0267462 with a tryptophan to alanine substitution at amino acid position 620 | | | pDM317 (Veltman et al., 2009) |
| GFP-Dd5P4^D319G^ | DDB_G0267462 with an aspartate to glutamine substitution at amino acid 319 | | | pDM317 (Veltman et al., 2009) |
| GxcU-GFP | GxcU (*D. intermedium*) | | | pDM323 (Veltman et al., 2009) |
| OIBP-GFP | DDB_G0275301 | | | pDM323 (Veltman et al., 2009) |
| PIO-GFP | DDB_G0280015 | | | pDM323 (Veltman et al., 2009) |

Loovers, H.M., Kortholt, A., de Groote, H., Whitty, L., Nussbaum, R.L., and van Haastert, P.J. (2007). Regulation of phagocytosis in Dictyostelium by the inositol 5-phosphatase OCRL homolog Dd5P4. Traffic *8*, 618-628.

Loovers, H.M., Veenstra, K., Snippe, H., Pesesse, X., Erneux, C., and van Haastert, P.J. (2003). A diverse family of inositol 5-phosphatases playing a role in growth and development in Dictyostelium discoideum. The Journal of biological chemistry *278*, 5652-5658.

Veltman, D.M., Akar, G., Bosgraaf, L., and Van Haastert, P.J. (2009). A new set of small, extrachromosomal expression vectors for Dictyostelium discoideum. Plasmid *61*, 110-118.
