## Supplementary Figure 1 for "Lowe Syndrome-linked endocytic adaptors direct membrane cycling kinetics with OCRL in *Dictyostelium discoideum*"

**A**

**B**

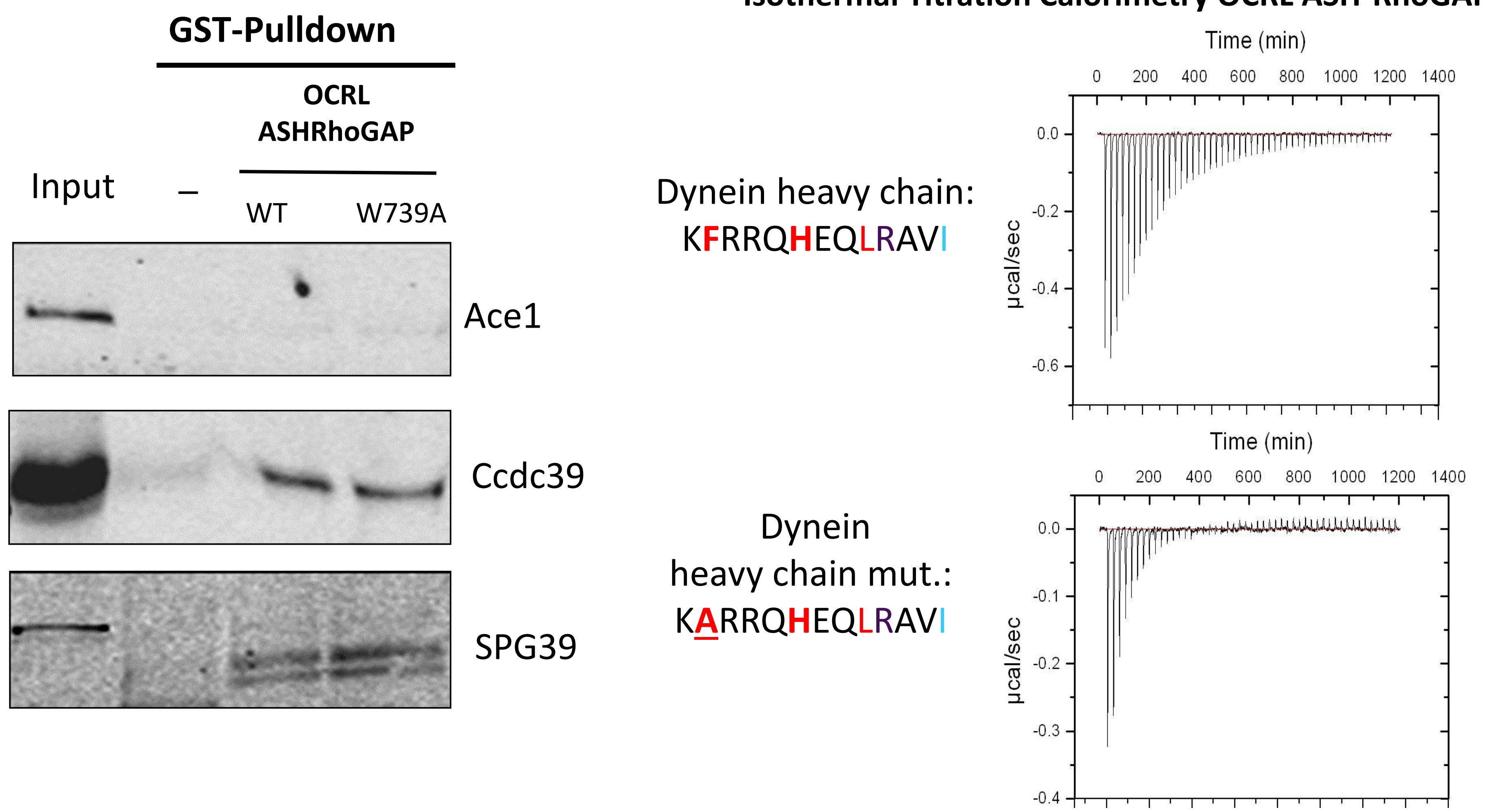


**C**

| Peptide | Protein | M.S(*) | GST-PD | W.B | Array | ITC(+) |
|---|---|---|---|---|---|---|
| **Human proteins tested** | | | | | | |
| **CFSTLHDWYGQEI** | **hSes2** | **Y** | **Y** | **Y(+)** | **Y** | **–** |
| **PFARLHECYGQEI** | **hSes1** | **Y** | **Y** | **Y(+)** | **Y** | **Y(+)** |
| **SFQQRHESLYRP** | **hAPPL1** | **N** | **Y** | **Y(+)** | **Y** | **Y(+)** |
| **AFHQRHETFRNAI** | **Frabin** | **N** | **Y** | **Y** | **Y** | **–** |
| IFHLYHKKTLFYS | FGD5 | N | N | **–** | **–** | **–** |
| VFQLLHKNMLFYV | FGD6 | N | N | **–** | **–** | **–** |
| DFIMMHCVFMPNT | SPG39 | N | – | **N** | **Y** | **–** |
| LFSIRHRSLHRHS | Ace1 | N | – | **N** | **Y** | **–** |
| DFRKIHNERQELI | Ccdc39 | N | – | **N** | **Y** | **–** |
| VFSQLHELRQQDL | WDR81 | N | – | **–** | **Y** | **–** |
| RFRKLHCTRNFIH | PACR | N | – | **–** | **Y** | **–** |
| KFRRQHEQLRAVI | **Dynein.h.c** | N | – | – | N | N |
| PFARLAECYGQEI | **hSes1(mut)** | – | N(+) | N(+) | N | N(+) |
| **Dictyostelium proteins tested** | | | | | | |
| **DFSSQHQQYSFDQ** | **OIBP** | **Y** | **Y** | | | |
| **DFLTLHKAYIDEW** | **PIO** | **Y** | **Y** | | | |
| **SFIATHTAYNKEN** | **GxcU** (#2) | **Y** | **Y** | | | |
| SFQLIHPIKSFTL | GxcU (#1) | Y | N | | | |
| GFVYIHCNLGLFK | Hwr (#1) | Y | N | | | |
| CFGTTHFCDTCHD | Hwr (#2) | Y | N | | | |
