## Supplementary figures and images for "Lowe Syndrome-linked endocytic adaptors direct membrane cycling kinetics with OCRL in *Dictyostelium discoideum*"

### Supplementary Figure 2

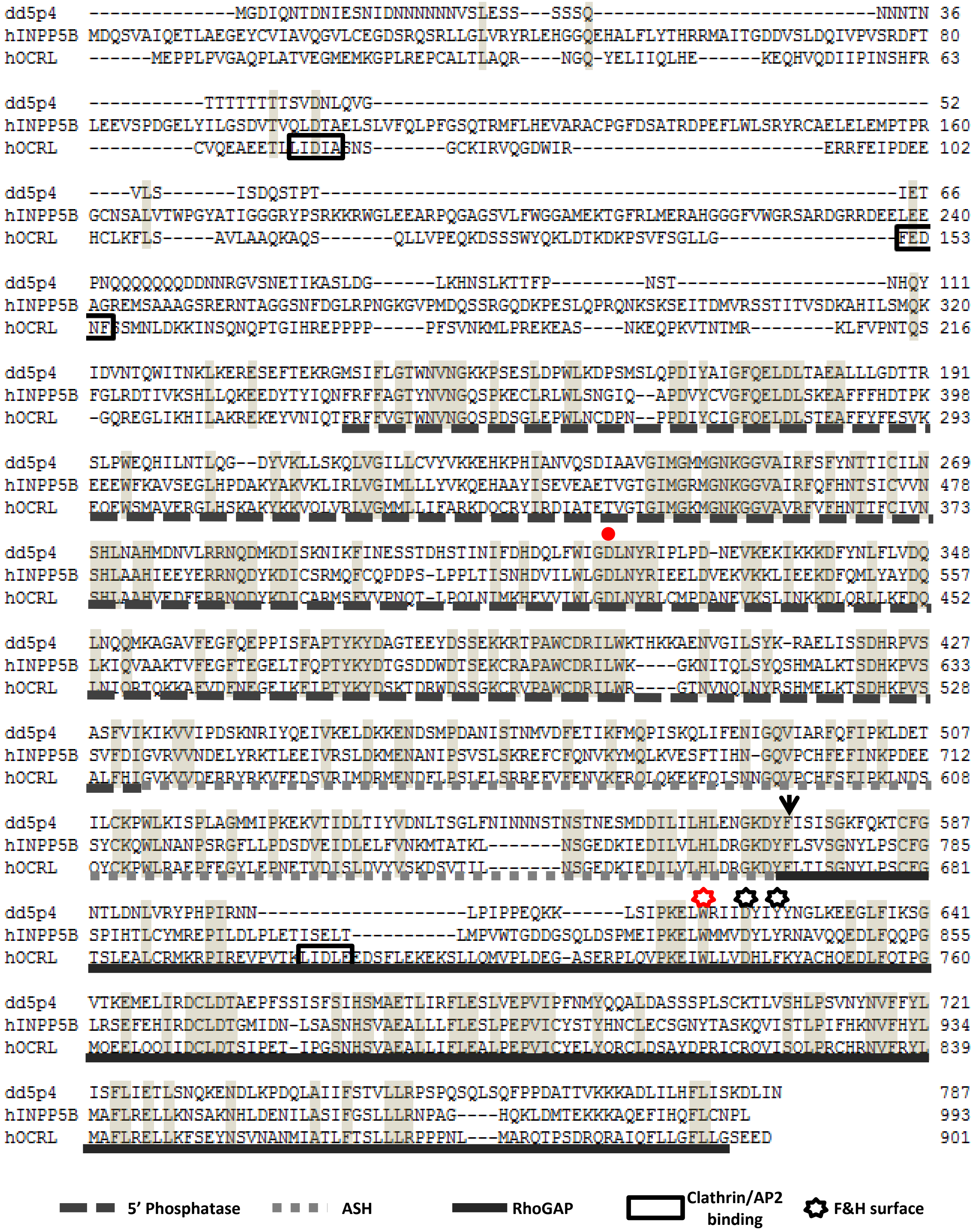

### Supplementary Figure 3

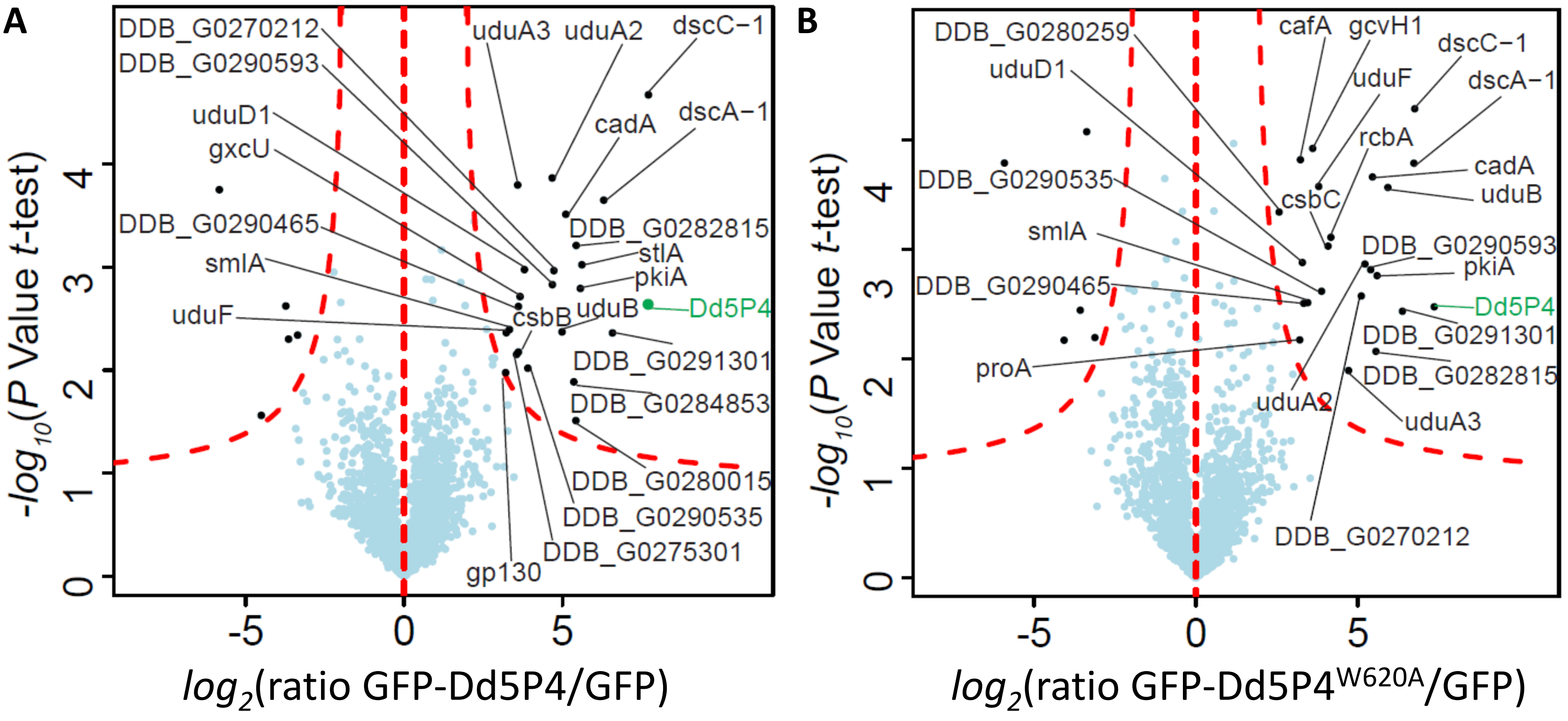

### Supplementary Figure 4

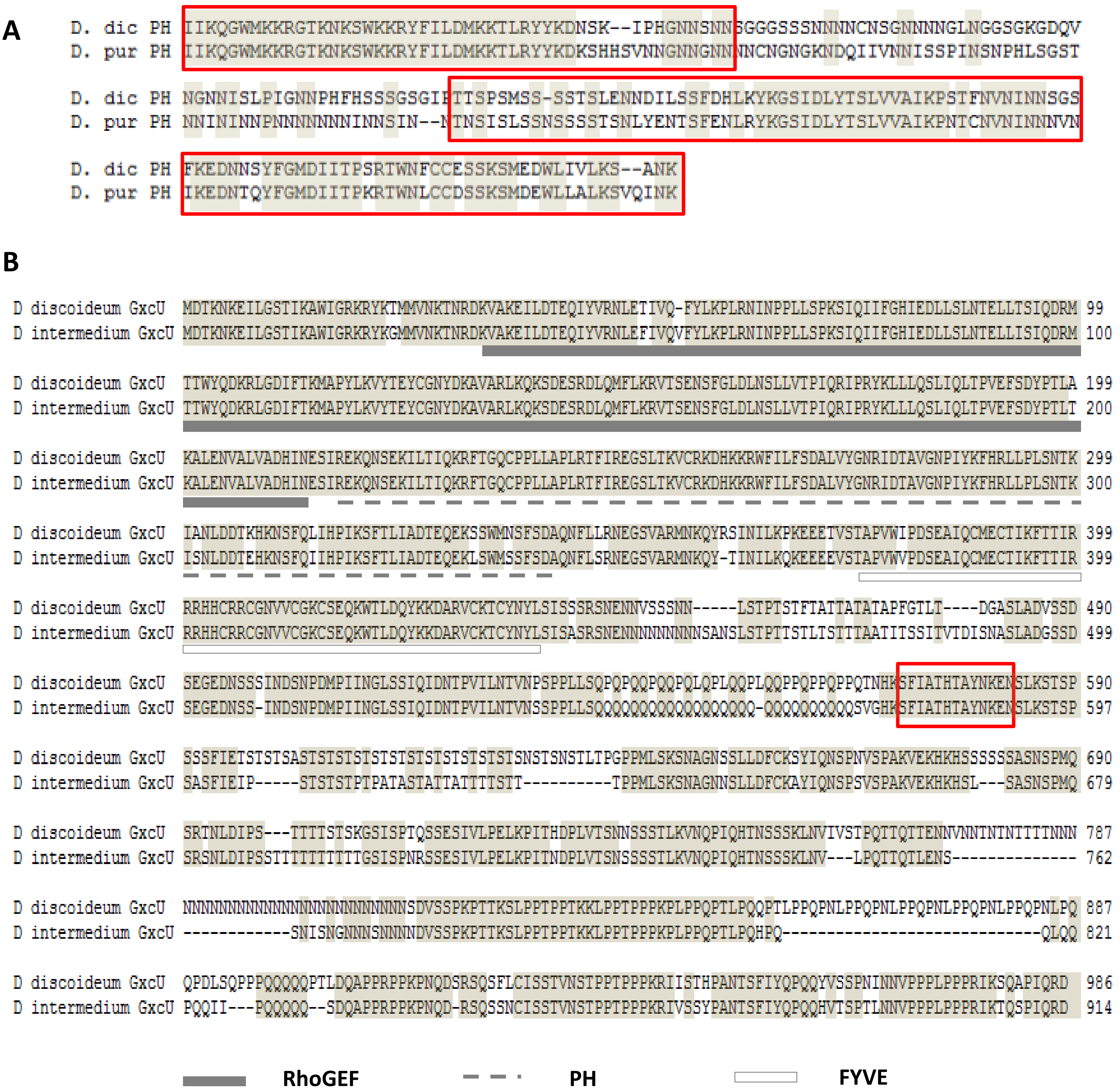

### Supplementary Figure 5

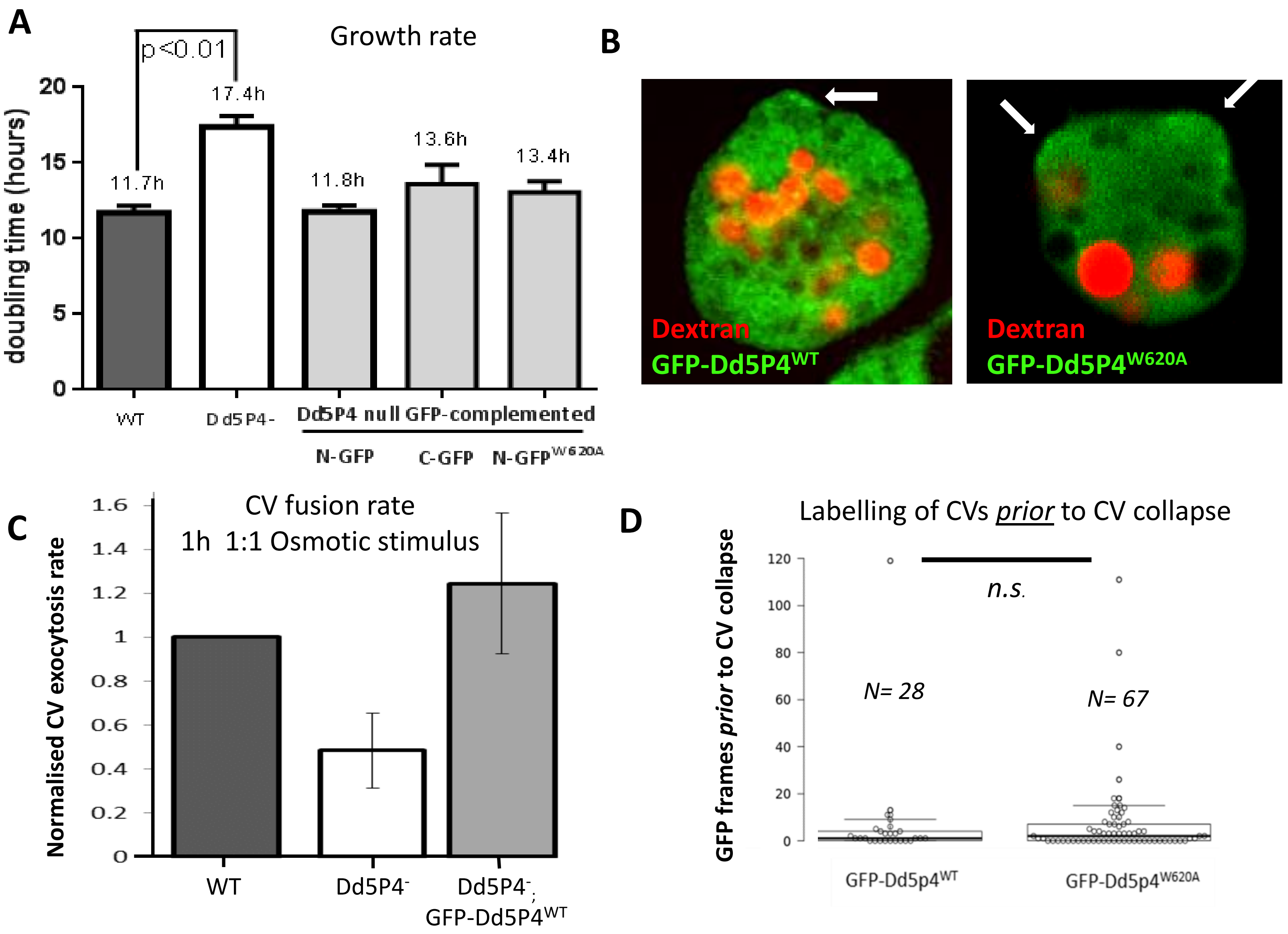
